# The growth hormone gene has evolved independently in *African Pygmy Mouse Mus minutoides*

**DOI:** 10.1101/2021.04.12.439429

**Authors:** Sumito Matsuya, Hiroyuki Imai, Yasuo Kiso, Ken Takeshi Kusakabe, Kiyoshi Kano

## Abstract

*Mus minutoides* (the African pygmy mouse) is one of the smallest mammals. We determined the nucleotide sequence of the *growth hormone* (*Gh*) gene and the sequence of the putative coding region in *M. minutoides*, where is predicted to be distinct in the functional and transcriptional regulatory regions between *M. minutoides* and *Mus musculus* (the House mouse). To investigate the evolutionary characteristics of *Gh* in *M. minutoides*, we constructed a phylogenetic tree based on the putative amino acid sequences of *Gh* in *M. musculus* and mammals by neighbor-joining method, suggesting that *Gh* diverged relatively earlier than other *Mus* genus and may have evolved independently in *M. minutoides*. Furthermore, analysis of *Gh* gene expression levels showed a tendency to be higher in *M. minutoides* than in *M. musculus*. Our results suggest that *Gh* may have evolved independently in *M. minutoides* and may have different functions and signaling in *Mus* genus.

---

Mammals have various body sizes, ranging from large whales to small rodents. Each species also has an inherent size, but it is not clear how this is determined and maintained. *Mus minutoides* (African pygmy mouse) is one of the smallest mammals in the world, measuring about 30 mm in body length and about 3 grams in body weight. *M. minutoides* originally inhabit the south of Saharan in Africa and is now bred as a pet animal in many countries [6, 11]. The sexual maturity of *M. minutoides* is about 8 weeks old, the gestation period is about 20 days, and the lifespan is about 2 years [17], which is similar to that of the common laboratory mice, but the chromosomes of *M. minutoides* have unique characteristics that differ from those of *Mus musuclus. M. minutoides* can be broadly divided into two major groups: the Southern Clade, which is widely distributed in South Africa, and the Eastern Clade, which is restricted to the eastern part of the South Africa region. Chromosome numbers are 2n=18, 32, 34 in the Southern Clade, but not have been identified in the Eastern Clade [6].

Interestingly, in addition to the X and Y chromosomes, *M. minutoides* has a unique chromosome called X*, which is shorter than the original X chromosome and has a special sex-determination pattern [13]. X chromosomes inactivation and sex determination, male hormones, and characteristics of X*Y female have been well studied [9, 14, 16]; however *M. minutoides* is one of the smallest mammals, despite its obvious characteristics, few studies have focused on its size.

Many factors seem to be involved in determination of unique body size and growth rates in different animal species, and they are thought to interact in complex manner. One of the most representative factors involved in growth is growth hormone (Gh). Many studies have been conducted to identify the structure, functional and binding sites of Gh and Gh receptors, and to determine the regulatory mechanisms of growth and homeostasis [1, 4, 7, 10, 18]. In animals, Gh has been also well-studied, and the sequences of the *Gh* gene have been identified not only in rodents but also in dogs [2], cats [5], and pandas [12] and so on. In general, dwarfism is also due to impaired function of growth hormone regulators in the anterior pituitary gland and neuroendocrine and tissue transcription factors in the anterior pituitary in mice [3, 15].

In addition, understanding the regulatory mechanisms of Gh and other pituitary hormones is particularly important in body size determination mechanisms because many spontaneous dwarf mice may be deficient in pituitary hormones.

The purpose of this study is to elucidate the basic mechanisms that determine the size of *M. minutoides* by analyzing firstly the nucleotide sequences of Gh, which is involved in animal body size. The results indicated that growth hormone evolved independently within the *Mus* genus at the presumed amino acid sequence level, suggesting that it may be responsible for the characteristic small body size of *M. minutoides*.

All animal experiments were approved by the Experimental Animal Care and Use Committee of Yamaguchi University (protocol number: 291). *Mus minutoides* used in this study were prepared for experiments immediately after purchase (Increase, Inc., Himeji, Japan). *Mus musculus* (C57BL/6J) were purchased from Japan SLC (Hamamatsu, Japan) and housed in groups with *ad libitum* access to water and food. Primers were designed based on the nucleotide sequence of the Ensembl database (Mouse (GRCm38.p6)) of the *growth hormone* (*Gh*) gene, and the genomic DNA extracted from the tail of *Mus minutoides* was used for sequencing. The primers used for nucleotide sequencing were designed to divide the entire gene length into three regions based on the sequence of *M. musculus*, and the sequence of the putative genomic region of the *Gh* gene in *M. minutoides* was determined. The primer sequences are listed in Table 1. Genomic PCR was performed and sequenced to search for differences between *M. musculus* and genes that may have a significant effect on body size determination. The *Gh* gene was amplified by PCR using the genome extracted from *M. minutoides* and compared with *M. musculus*. The *Gh* gene of *M. musculus* consists of a 1695 bp sequence and contains five exons. Alignment of the nucleotide sequence in the Ensembl database of *M. musculus* showed substitutions, deletions, and insertions in all regions including exons in *M. minutoides* (Fig. 1A). We have already registered the DNA sequence of the *Gh* gene of *M. minutoides* to DNA Data Bank of Japan (accession number: LC5671320).

**Fig. 1.**
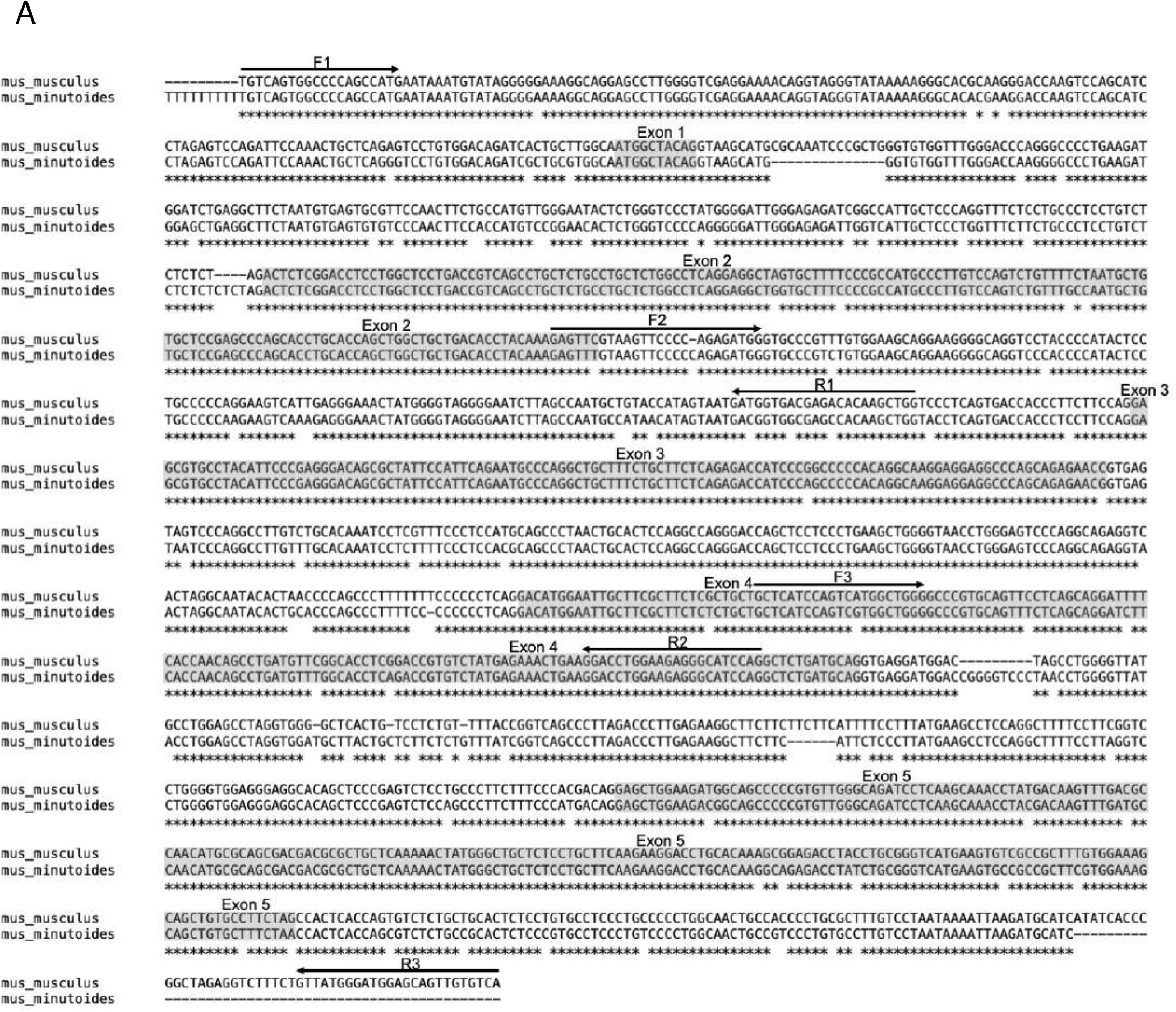
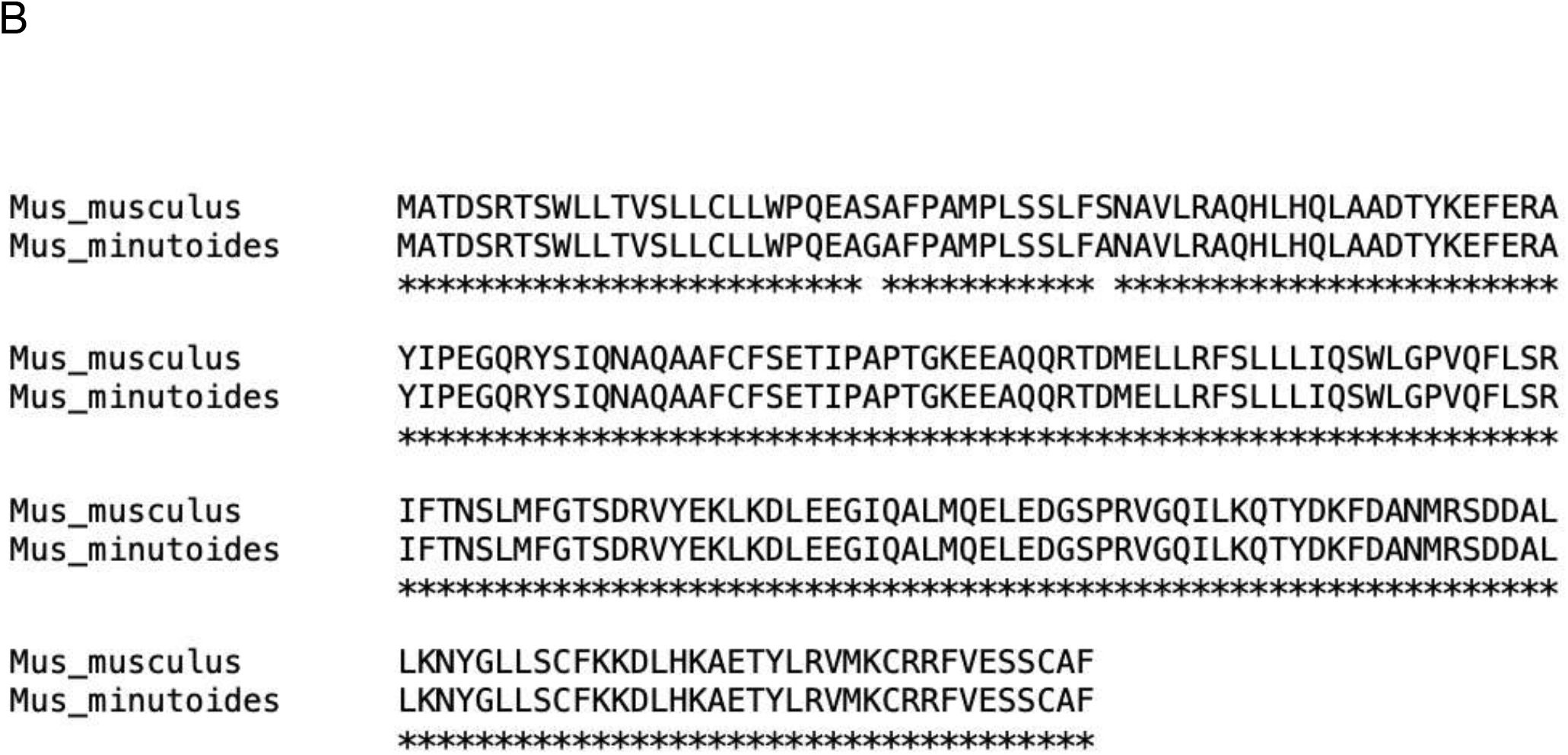
Alignment of a putative amino acid sequence of Gh between *M. minutoides* and *M. musculus*. The amino acid sequence of growth hormone was presumed by a nucleic acid sequence of *M. minutoides*. The 25th and 37th amino acid sequences of *M. musculus* are serine, respectively, while two sites are different in *M. minutoides*: glycine and alanine.

Furthermore, the sequence of *M. minutoides of* the *Gh* gene was translated into a putative amino acid sequence and compared with the amino acid sequence of *M. musculus* using BLAST. The length of the putative amino acid sequence was 216 amino acids in *M. minutoides*, and identical to that of *M. musculus*. At the DNA sequence level, the *Gh* gene in *M. minutoides* had many sequences that differed from those in *M. musculus*, but the homology at the putative amino acid level was very high except for two amino acids, suggesting that *M. minutoides* also may have a functional Gh. The results showed that only the 25th and 37th amino acid sequences of *M. musculus* are serine, respectively, while two sites are different in *M. minutoides*: glycine and alanine (Fig. 1B). These mutations were found in the N-terminal region responsible for signal transduction and in the region that promotes GH protein maturation, respectively [8]. The 25th amino acid in *M. musculus* is serine, while the 25th amino acid in *M. minutoides* is glycine, suggesting that the binding ability of GH protein to receptors may be altered. Serine, the 37th amino acid in *M. musculu*s, is part of the site responsible for the maturation of the GH protein, suggesting that the maturation process or post-translational modification of the GH protein in *M. minutoides* may be different from that in *M. musculus*.

Next, to examine whether these amino acid differences affect Gh gene expression, Gh gene expression in the pituitary gland of adult male *M. minutoides* (n=3). Table 2 lists the primers used to detect *Gapdh and Gh. M. musculus* was examined using qRT-PCR and found that the expression of the *Gh* gene was higher in *M. minutoides* than in *M. musculus* (Fig. 2). It is unlikely that the sequence of the *Gh* gene is directly related to the secretion of Gh protein, but the relatively high expression of the *Gh* gene in *M. minutoides* may not be directly related to body size. In dogs, growth promotion has been correlated with the level of Igf1, a downstream factor of GH, rather than with the GH secretory [8]. In our preliminary experiments, we have found that the expression level of the *Igf1* gene in *M. minutoides* is much lower than that in *M. musculus*. Although the levels of Gh and Igf1 in the serum need to be measured, the relatively high expression of the *Gh* gene in *M. minutoides* may be due to negative feedback of Igf1 levels.

**Fig. 2.**
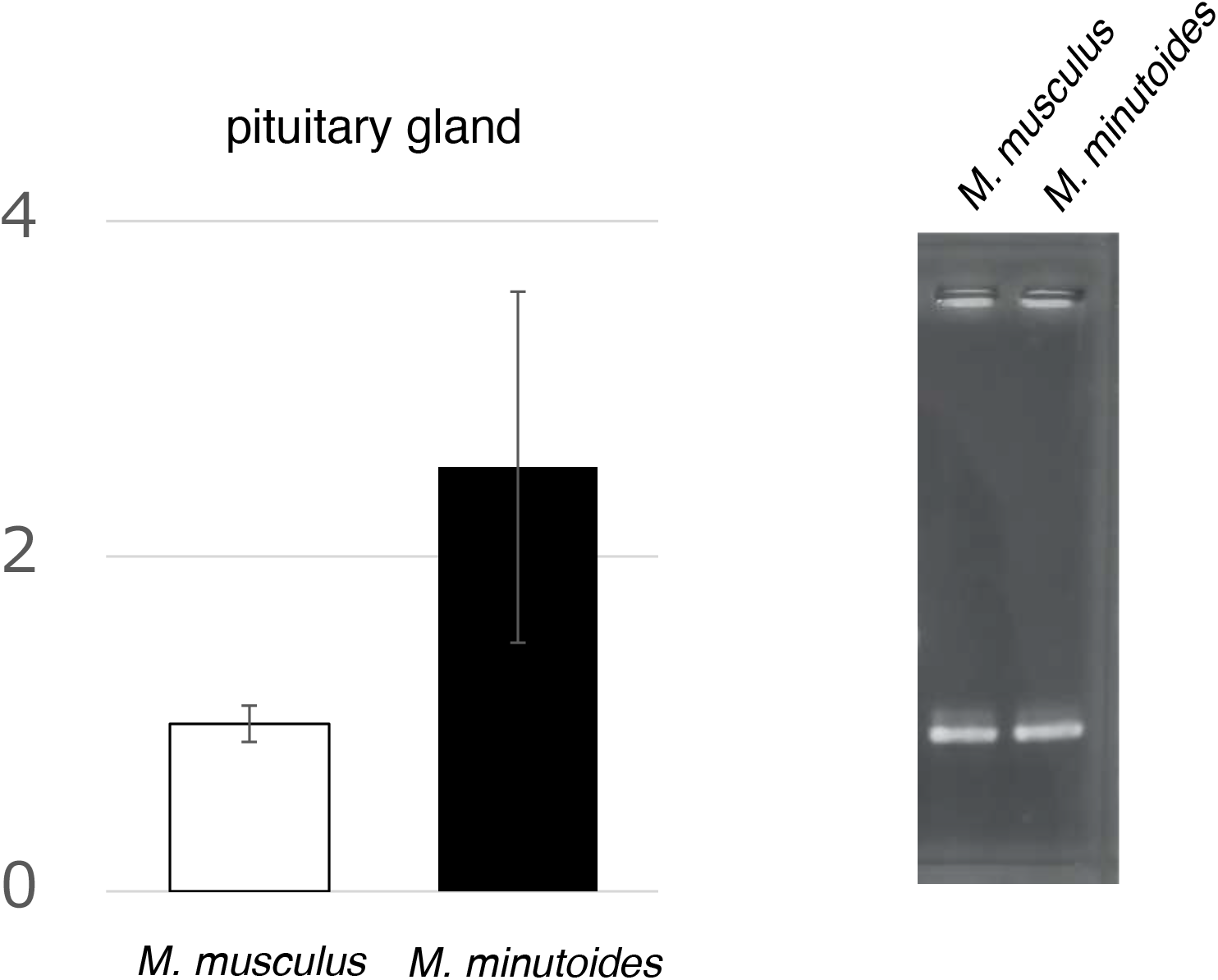
Transcript levels of *Gh* in the pituitary gland of *M. minutoides* and *M. musculus*. Left: The relative expression level for *Gh* was normalized against the *GAPDH* expression level. The expression of the Gh gene in *M. minutoides* was higher than those in *M. musculus*. Data represent the mean ± SD. Right: RT-PCR analysis was performed for confirmation of the specificity of the primer set.

Finally, to study the evolutionary characteristics of the amino acid sequence of Gh in *Mus minutoides*, we constructed a phylogenetic tree of Gh in other *Mus* genus and mammals (Fig. 3) by neighbor-joining method using the Poisson model by MEGAX with 1,000 bootstraps (The Biodesign Institute, Tanpe, AZ, USA), suggesting that the Gh diverged relatively earlier than in the other *Mus* genus and may have evolved independently in *M. minutoides*. This indicates that *M. minutoides* might have a unique growth system of determining body size through mediated Gh hormone.

**Fig. 3.**
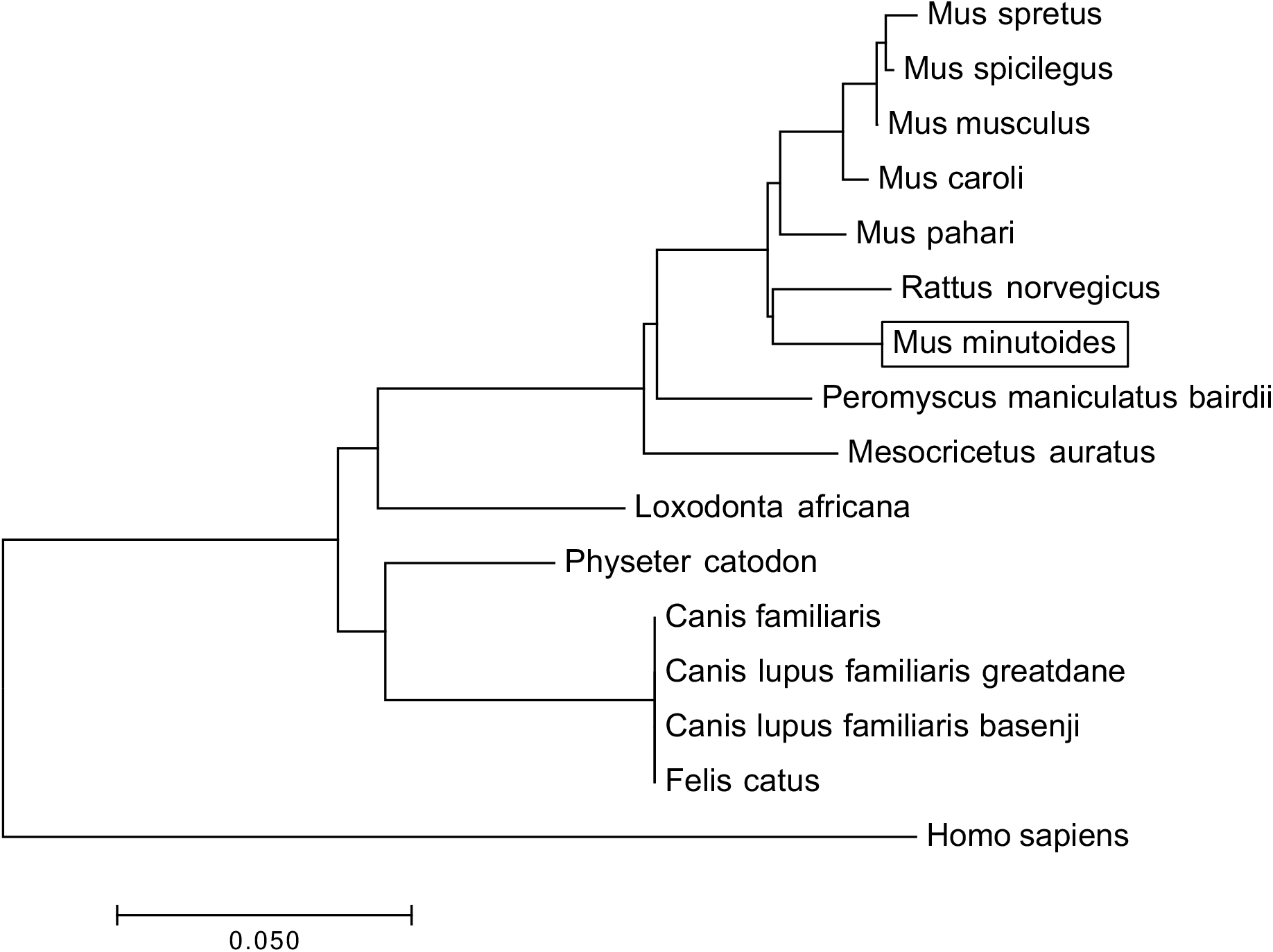
Phylogenetic tree of the vertebrate growth hormone (Gh). The tree was constructed with the maximum likelihood method using MEGAX from *M. minutoides* and 16 other vertebrate Gh amino acid sequences. Gh amino acid sequences were obtained from the Ensembl database.

The present study suggests that *M. minutoides* evolved independently in Gh and may have different functions and signaling from *M. musculus*. In order to further elucidate the characteristic growth-related characteristics of *M. minutoides*, particularly Gh, it will be necessary to analyze related factors such as growth hormone receptors and Igf1.

## Acknowledgments

The authors thank the ARCLAS animal facility and the technical expertise of The DNA Core facility of the Center for Gene Research, Yamaguchi University. We also thank W. Fujii for his comments on the research. We are grateful to anonymous reviewers for their insightful comments.

## Figure legends

**Fig. 1** Alignment of a nucleic acid sequence of the genomic region and a putative amino acid sequence of *Gh* gene. (A) Alignment of a nucleic acid sequence of the genomic region of *Gh* gene between *M. minutoides* and *M. musculus*. Arrows indicate primers for the nucleotide sequence. The upper row is the sequence of *M. musculus* and the lower row is that of *M. minutoides*. * indicates matching nucleic acids between *M. minutoides* and *M. musculus*. Deletions and insertions across the entire region, including exons, was observed in the genomic DNA of *M. minutoides*. The shaded indicates five exons of *M. musculus*. (B) Alignment of a putative amino acid sequence of Gh between *M. musculus* and *M. minutoides*. The amino acid sequence of growth hormone was presumed by a nucleic acid sequence of *M. minutoides*. The 25th and 37th amino acid sequences of *M. musculus* are serine, respectively, while two sites are different in *M. minutoides*: glycine and alanine.

**Fig. 2** Transcript levels of *Gh* in the pituitary gland of *M. minutoides* and *M. musculus*. Left: The relative expression level for *Gh* was normalized against the *Gapdh* expression level. The expression of the *Gh* gene in *M. minutoides* was higher than those in *M. musculus*. Data represent the mean ± SD. Right: RT-PCR analysis was performed for confirmation of the specificity of the primer set.

**Fig. 3** Phylogenetic tree of the vertebrate growth hormone (Gh). The tree was constructed with the maximum likelihood method using MEGAX from *M. minutoides* and 16 other vertebrate Gh amino acid sequences. Gh amino acid sequences were obtained from the Ensembl database.

---

**Table S1.**
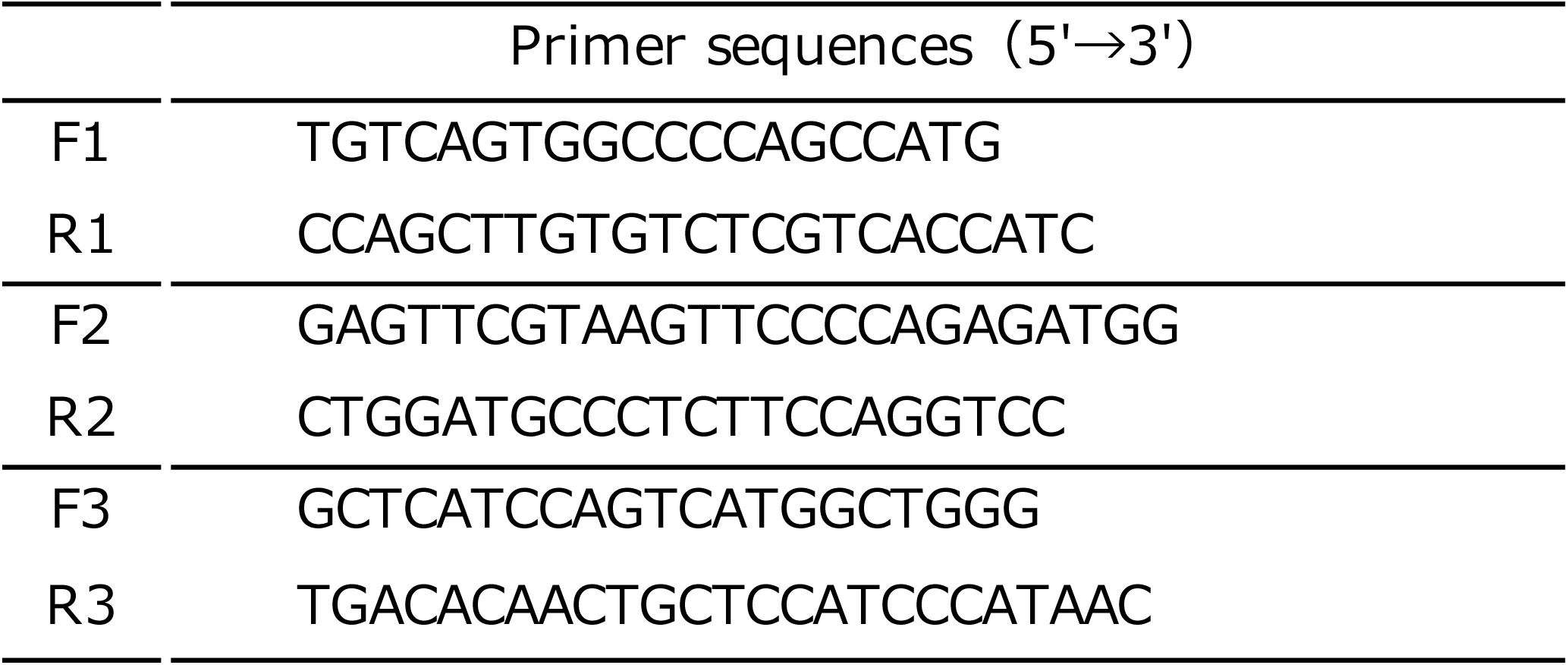
Primer sequences for the genome sequence of Gh

**Table S2.**
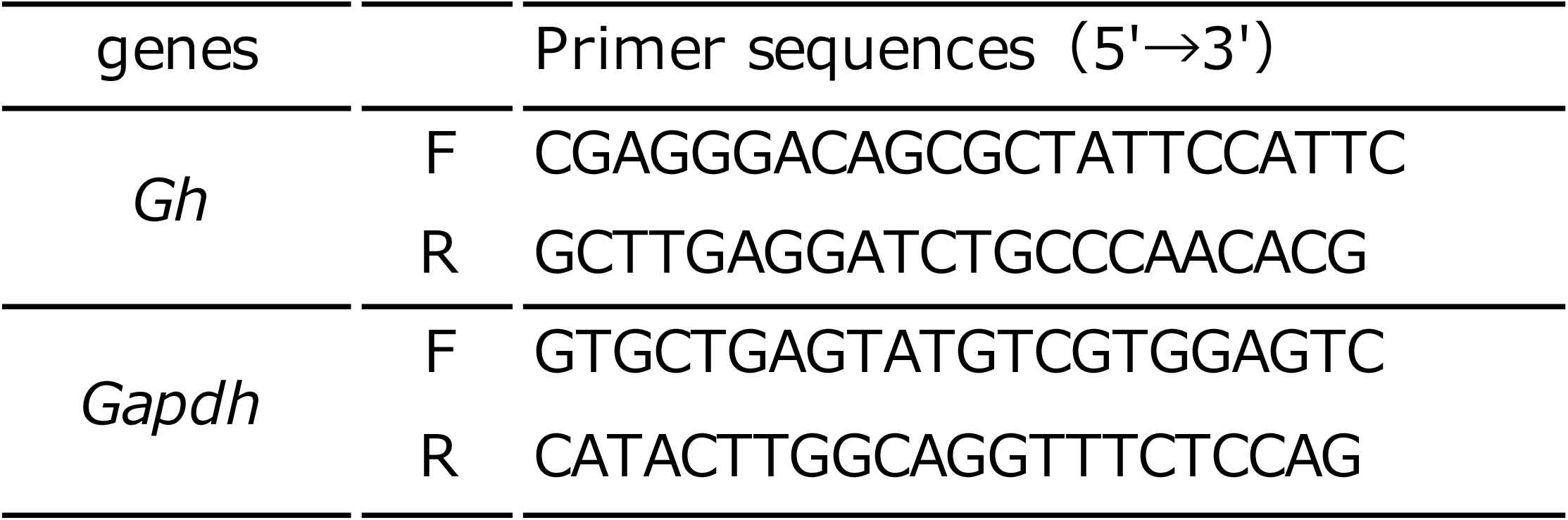
Primer sequences for RT-PCR

